# A SOD1-dependent mitotic DNA damage checkpoint

**DOI:** 10.1101/2022.10.26.513831

**Authors:** Rachel Gatenby, Nan Li, Priya Lata, Thomas Walne, Afzaal Tufail, Alexander Breitweiser, Ruth Thompson

**Author notes:** These authors contributed equally to this work (Rachel Gatenby, Nan Li).

## Abstract

In the event of DNA damage, the cell cycle can be slowed or halted to allow for DNA repair. The mechanisms by which this occurs are well-characterised in interphase, although the mechanisms underpinning mitosis slowing in response to damage are unclear. Canonical checkpoints and DNA repair pathways are largely repressed in mitosis, and whilst there is some level of mitotic DNA synthesis and repair, the bulk of DNA damage is processed for post-mitotic repair. How the decision is made between mitotic DNA repair and post-mitotic DNA repair is not known. We have identified the antioxidant enzyme Superoxide Dismutase 1 (SOD1) as an essential factor mediating delayed mitotic progression in response to DNA damage and replication stress. Cells depleted of SOD1 no longer exhibit DNA damage dependent mitotic delay, and display increased levels of damaged centromeres and mitotic defects. Whilst reactive oxygen species (ROS)-inducing agents also lead to SOD1-dependent mitotic delay, intracellular ROS levels do not correlate with mitotic arrest. SOD1 appears to play an important role in DNA repair in interphase and is recruited to the nucleus in response to DNA damage. In addition to control of mitotic progression in response to genotoxic stress, SOD1 also plays a major role in mitotic DNA synthesis. SOD- depleted cells show reduced levels of mitotic EdU incorporation in response to either replication stress or DNA breaks, seemingly in tandem with Rad51 andSOD1-depletion induced mitotic progression in the presence of DNA breaks is Rad52-dependent. We suggest that there are two responses to DNA breaks in mitosis; either arrest and mitotic repair or progression and post-mitotic repair; and these two pathways exist in a fine balance, controlled by a signaling cascade involving SOD1.

## Introduction

It is vital that cells maintain genomic integrity in order to pass on a faithful copy of their genetic material to the next generation. All the cells in our body are continuously exposed to genotoxic threats, with tens of thousands of DNA-damaging events occurring daily in each cell. The cellular response to DNA damage involves careful coordination of cell cycle control, DNA repair and programmed cell death.

In response to DNA damage, the phosphatidylinositol 3-kinase-related kinases Ataxia Telangiectasia Mutated (ATM) and Ataxia Telangiectasia Rad3-Related (ATR) activate cell cycle checkpoints throughout interphase, resulting in cell cycle arrest at the G1/ S and G2/M boundaries, and slowing of DNA replication via Cyclin dependent kinase (CDK) inhibition [1]. Whilst cell cycle control and activation of DNA repair are well characterised in interphase, how the cell cycle responds to DNA breaks in mitotic cells remains unclear. Whilst we and others have reported slowed mitotic transit in response to DNA damage [2–5], it is generally accepted that there is no mitotic DNA damage-induced cell cycle checkpoint [6]. The reasoning for this is two-fold; Firstly, the interphase checkpoints all act via inhibition of CDKs necessary for cell cycle progression [7], and several mechanisms prevent the initiation of such mechanisms during mitosis, reviewed in [6]. Low levels of transcription in mitosis prevent CDK inhibition via p21 induction, and furthermore the mitotic kinase Plk1 acts to inhibit Wee1 and, both of which are required for the inhibitory phosphorylation of CDK1 at Tyr 15 [6]. Secondly, the canonical DNA damage response (DDR) is largely inhibited in mitosis to avoid the risk of telomere fusion [8], which has led to the hypothesis that there is no requirement for a mitotic DNA damage checkpoint. Whilst the signalling cascade which responds to DNA double strand breaks (DSB) in interphase can be initiated in mitosis, the cascade is thought to be attenuated in mitotic cells, to be reinstated following the completion of mitosis [9]. Studies in fly [10] and human [11] cells suggest that broken chromosomes can be “tethered” together in mitosis to allow for faithful segregation of potential acentric chromosomes.

Genomic instability - sometimes resulting in full chromosome rearrangements - is a hallmark of cancer. Many chromosome rearrangements occur at the same sites in chromosomes, known as “common fragile sites” or CFS [12,13]. CFS represent difficult-to-replicate areas of the genome which usually replicate late in S phase and in some cases in early G2 [14] and mitosis [15] in response to replication stress. In mitosis, the endonucleases MUS81/EME1 are recruited to CFS and induce DNA breaks, whereupon the nuclease activity of MUS81 promotes POLD3-dependent Mitotic DNA synthesis (MiDAS) to minimise chromosome mis-segregation in anaphase [15]. Two types of MiDAS have been observed; one which occurs at sites of DNA DSB and is Rad52 dependent: termed DDR-associated MiDAS, and another which occurs in the absence of DNA DSB and is dependent on Rad51: termed non-DDR-associated MiDAS [16]. Mitotic DNA synthesis also occurs at sites of laser-induced DNA DSB [17], indicating that this phenomenon is not restricted to sites of DNA under-replication.

Superoxide Dismutase 1 (SOD1) has a well-established role as a dismutase of toxic superoxide radicals (O_2_-), which it converts to the more stable and less toxic hydrogen peroxide and dioxygen [18]. More recently, SOD1 has been implicated in the DDR with elevated levels of DNA damage observed in SOD1 mutant ALS cells [19,20] and furthermore over-expression of SOD1 leads to activation of the DDR in SOD1 mutant cells [21]. Loss of SOD1 has also been shown to confer sensitivity to DNA damaging agents and lead to downregulation of the ATM pathway in yeast [22]. In addition, SOD1 has also been implicated in regulation of gene expression. SOD1 is also reported to be activated by the DDR factors ATM and Chk2 [23–25] and in turn, to act as a transcription factor to initiate expression of ROS-reducing and DNA damage-related genes [23]. In this manuscript, we further characterise the role of SOD1 in DNA repair and demonstrate a novel role for SOD1 in the control of mitotic progression in response to DNA damage.

## Experimental procedures

### Cell Culture and Reagents

HeLa, MCF7 and MRC5 cells (ATCC) were cultured as previously described [5]. Irradiation was carried out using the CIB/IBL 437 Cesium-137 irradiator. Where indicated, cells were treated with Carboplatin (Sigma-Aldrich), Okadaic acid (Cell Signaling), Sodium Selenate (Sigma-Aldrich), Temozolomide (Sigma-Aldrich), H_2_O_2_ (EMD Millipore), Pyocyanin (Sigma Aldrich), Alisertib (Selleckchem), aphidicolin (Santa Cruz), RO3306 (Cayman Chemical), B02 (Sigma-Aldrich), AICAR (Sigma-Aldrich) at indicated concentrations and with indicated length of period. Antibodies against SOD1, BubR1 Rad51, Mad2 and Securin were obtained from Abcam, Bub3, Cdc20 and Myc-Tag were from Cell Signaling, Actin and CENP B were from Santa Cruz and GPx1 was from Novus Bio. Appropriate secondary antibodies conjugated to horseradish peroxidase were used (DAKO).

### RNAi and DNA transfection

siRNA transfections were performed using Dharmafect 1 siRNA transfection reagent (Horizon), DNA transfections using Lipofectamine 2000 (Thermo-Fisher Scientific) and siRNA/ DNA contranfections using Dharmafect Duo (Horizon) according to the manufacturer’s instructions. Cells were treated 48 hours post-transfection. siControl, siBubR1, siATM, siATR, siChk1, siMre11, siRad50, siNBS1, siMDC1, siSOD1 siGenome SMARTpool siRNA pools were from Horizon Discovery.

single siRNA sequences used were:

siControl (UAAUGUAUUGGAACGCAUA)TT
siBubR1 (GATTTAGCACATTTACTAT)TT
siSOD1-5 (UCGUUUGGCUUGUGGUGUA)TT
siSOD1-7 (GUGCAGGGCAUCAUCAAUU)TT

PP2A-Myc-DDK construct in pCMV6 Origene (NM_002715, RC201334), SOD1 optimised construct was custom made by Eurofins and subcloned into pCMV6 with a Myc-DDK tag.

### Immunofluorescence Microscopy

HeLa cells (5 x 10^4^) were seeded directly onto coverslips fixed in methanol or paraformaldehyde, permeabilised in 0.2% Triton-X100, blocked in 5% BSA and stained with the indicated antibodies. Alexa-Fluor 488 and AlexaFluor 594 secondary antibodies (Invitrogen) were used. In the final wash, cells were incubated with DAPI (Life technologies) and mounted to slides using Immumount (Thermo Fisher Scientific).

### Live cell Imaging

48 hours post transfection with indicated siRNAs, cells were trypsinised, exposed to ionising radiation (IR) whilst in suspension and reseeded to 24 well plates. Once adhered to the plates (3-4 hours post IR), the cells were loaded into imaging system, or following chemical administration as indicated. Live cell images were captured using ZEISS Cell discoverer 7 microscope every 5 minutes for a duration of 20 hours.

### Intracellular ROS Assay

ROS levels were detected using the chloromethyl-2’,7’-dichlorodihydrofluorescein diacetate (CM-H2DCFDA) general oxidative stress indicator kit (Invitrogen) according to the manufacturer’s instructions. Fluorescence was measured at 495/ 527 nm on the SpectraMax M5e multi-mode microplate reader (Molecular Devices) and readings were normalised to the cell-free control.

### Comet Assay

100,000 cells per condition were isolated and the alkaline comet assay protocol was performed (Trevigen) according to the manufacturer’s instructions. Slides were imaged using the 10x/0.25 objective lens on the Eclipse TE200 fluorescent microscope (Nikon) using NIS Elements software (Nikon). The FITC channel was used to visualise the cells. Images were converted to the BNP file format in order to process using CometScore software. The average percentage tail DNA was calculated for 50 cells per condition and the overall mean determined across the experimental repeats.

### qRT-PCR

RNA extraction was performed using the manufacturer’s protocol and reagents from the RNeasy mini kit (Qiagen). The concentration (ng/μL) of the eluted RNA was quantified using the Nanodrop spectrophotometer (ND-1000). The RNA extracted underwent reverse transcription using the high-capacity RNA to cDNA kit (Applied Biosystems). qRT-PCR was conducted using Taqman probes according to the manufacturer’s instructions and processed on a 7900 RT-PCR machine using the SDS 2.4 software

### Mitotic EdU incorporation assay

MiDAS assay was performed 48 hours post transfection as described by Garribba et al [26]. Images were captured using Nikon ECLIPSE Ti2 confocal microscope.

## Results

### DNA-damage induced mitotic delay is dependent on SOD1

To investigate the mitotic cell cycle response to DNA damage, several cell lines were treated with DNA damaging agents followed by live cell time-lapse microscopy analysis. We observed that the average time spent in mitosis was significantly increased following the introduction of DNA damaging agents (**Figure 1A, Figure S1A**). This delay was not restricted to cells inflicted with DNA damage whilst in mitosis, as the delay can still be seen upwards of 16 hours post-irradiation, indicating that cells can escape earlier interphase checkpoints and enter mitosis with damaged DNA (**Figure 1B**). The mitotic delay was also expressed as an increase in cells expressing the mitotic marker protein phosphorylated Histone-H3, 16 hours after IR treatment (**Figure 1C**), which allowed us to screen for mitotic delay in the absence of selected DNA repair proteins. Unexpectedly, depletion of several DNA repair proteins previously implicated in mitotic progression (including ATM, ATR, Chk1, the MRN complex and MDC1) did not significantly reduce the mitotic population observed following treatment with IR, whereas cells depleted of the antioxidant protein SOD1 no longer exhibited mitotic delay following exposure to IR (**Figure 1D**).

**Figure 1.**
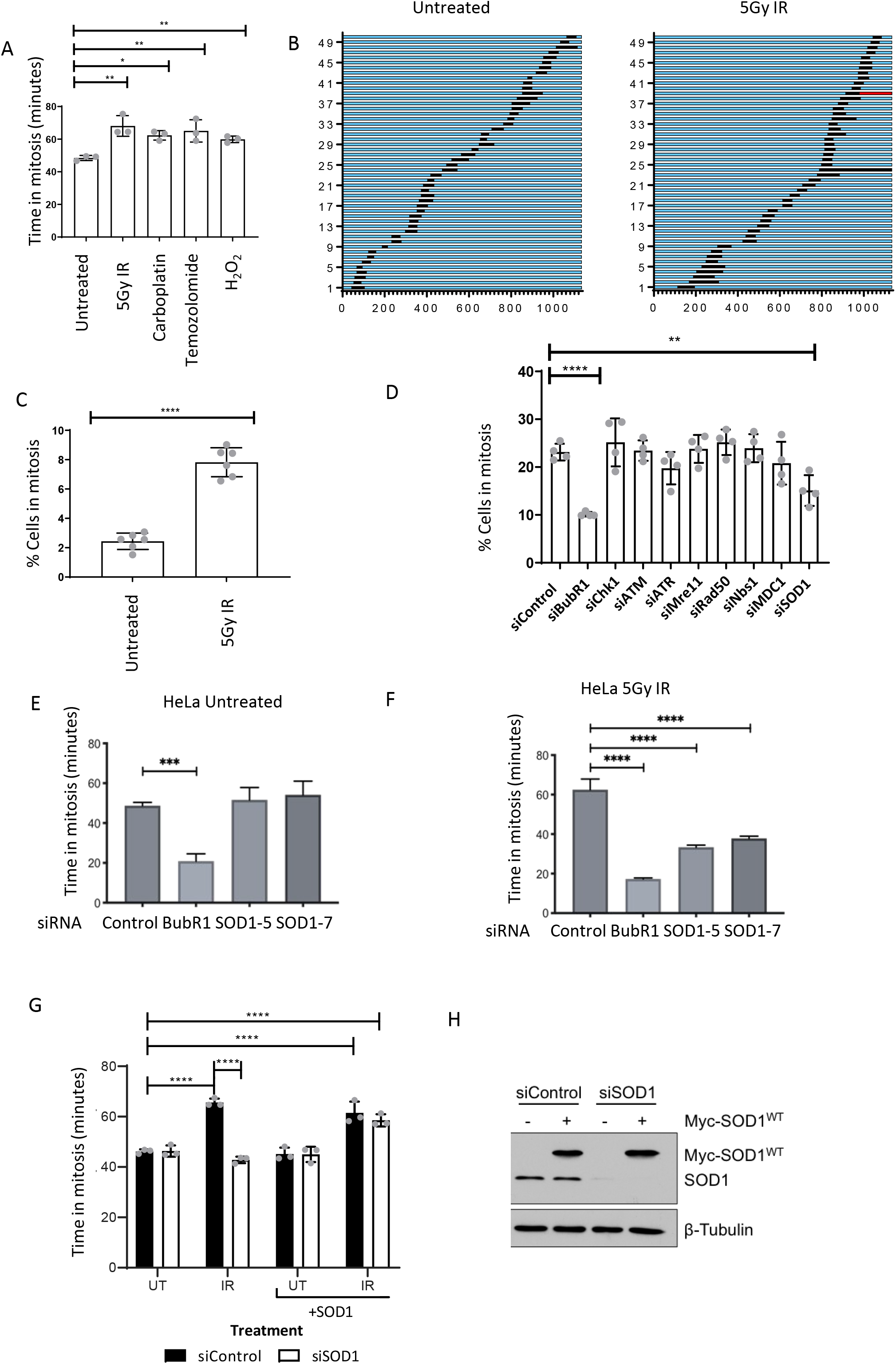
DNA damaging agents induce mitotic arrest which is dependent on SOD1. **A**. HeLa cells treated with the indicated agents were analysed by time lapse microscopy. Time in mitosis was scored from time cells rounded up in mitosis to time to cytokinesis. Error bars represent mean +/-SEM of 3 independent experiments (n>150). Results were analysed using a one-way ANOVA with Dunnett’s multiple comparison test. *p=0.05, **p=0.01, ***p=0.001, ****p=0.0001. **B**. 50 individual cells from a single repeat of A demonstrates mitotic delay persists for the duration of the experiment. **C.**Increase in mitotic population 16 hours post IR. Flow cytometry following pHH3 staining. Mean +/-SEM of 3 independent experiments. **D**. Cells incubated with the indicated siRNAs for 48 hours prior to treatment with 5Gy for 16 hours were stained for pHH3 and visualized by fluorescence microscopy. Error bars represent mean +/-SEM of 3 independent experiments (n>150). Results were analysed using a one-way ANOVA with Dunnett’s multiple comparison test. *p=0.05, **p=0.01, ***p=0.001, ****p=0.0001. **E & F**. HeLa cells incubated with the indicated siRNA were viewed by time lapse microscopy with and without IR treatment. Mean =/-SEM of 3 independent experiments (n>150). **G**. Ectopic expression of SOD1 rescues DNA damage induced mitotic delay following IR treatment. HeLa cells incubated with the indicated siRNA and cDNA were viewed by time lapse microscopy with and without IR treatment. Mean =/-SEM of 3 independent experiments (n>150). Results were analysed using a one-way ANOVA with Tukey post test. *p=0.05, **p=0.01, ***p=0.001, ****p=0.0001 **H.**Western blot demonstrating siRNA depletion and overexpression in g.

The siRNA used in the screen was based on pools of 4 individual siRNAs. Deconvoluted, all but one of the 4 SOD1 siRNAs used resulted in bypass of mitotic delay. The remaining siRNA yielded inconsistent data, and western blotting tests for SOD1 protein levels confirmed that it did not effectively reduce SOD1 protein levels (**Figure S1B and C**). We selected two siRNAs (5 and 7) for further study, and assessed the duration of mitosis in cells depleted of SOD1 or the Spindle Assembly Checkpoint (SAC) protein BUBR1 following exposure to IR Consistent with previous reports (REF), depletion of BUBR1 reduced mitotic transit time in unperturbed and IR treated cells alike, while depletion of SOD1 had no effect on mitotic progression in the absence of DNA damage (**Figure 1E**). However, when cells were treated with 5 Gy IR, SOD1-deficient cells no longer exhibited the increase in mitotic progression observed in cells transfected with a non-targeting control siRNA(**Figure 1F**), suggesting that SOD1 specifically regulates mitotic transit following DNA damage. The same result was observed in the breast cancer cell line MCF7 and, to a lesser extent, in the non-cancerous foetal lung cell line MRC5 (**Figure S1D and E**). Ectopic expression of siRNA-resistant SOD1 was sufficient to rescue DNA damage-induced arrest in SOD1-depleted cells, ruling out the possibility of off-target effects (**Figure 1G**).

### Mitotic delay is ROS independent

As the canonical function of SOD1 is the reduction of intracellular ROS, we investigated whether agents that increase ROS also lead to prolonged mitosis. We found that both hydrogen peroxide (H_2_O_2_) (**Figure 2A**) and pyocyanin (a ROS-inducing toxin produced by the bacterium *Pseudomonas aeruginosa*) (**Figure S2A)**induce SOD1-dependent mitotic delay to an extent similar to that observed in the presence of other DNA damaging agents (**Figure 2B**). SOD1 is required for the production of H_2_O_2_ from superoxide molecules so we initially theorized that the mitotic delay observed could be due to altered levels of cellular H_2_O_2_. However, the DNA damaging agents carboplatin and temozolomidealso induce SOD1-dependent mitotic arrest (**Figure 2B**), but do not produce significantly increased ROS (**Figure 2c**). Surprisingly, SOD1 siRNA depletion did not lead to a significant reduction of detectable H_2_O_2_ in any condition (**Figure 2C**), suggesting that altered ROS levels do not underpin SOD1 regulation of mitotic transit

**Figure 2.**
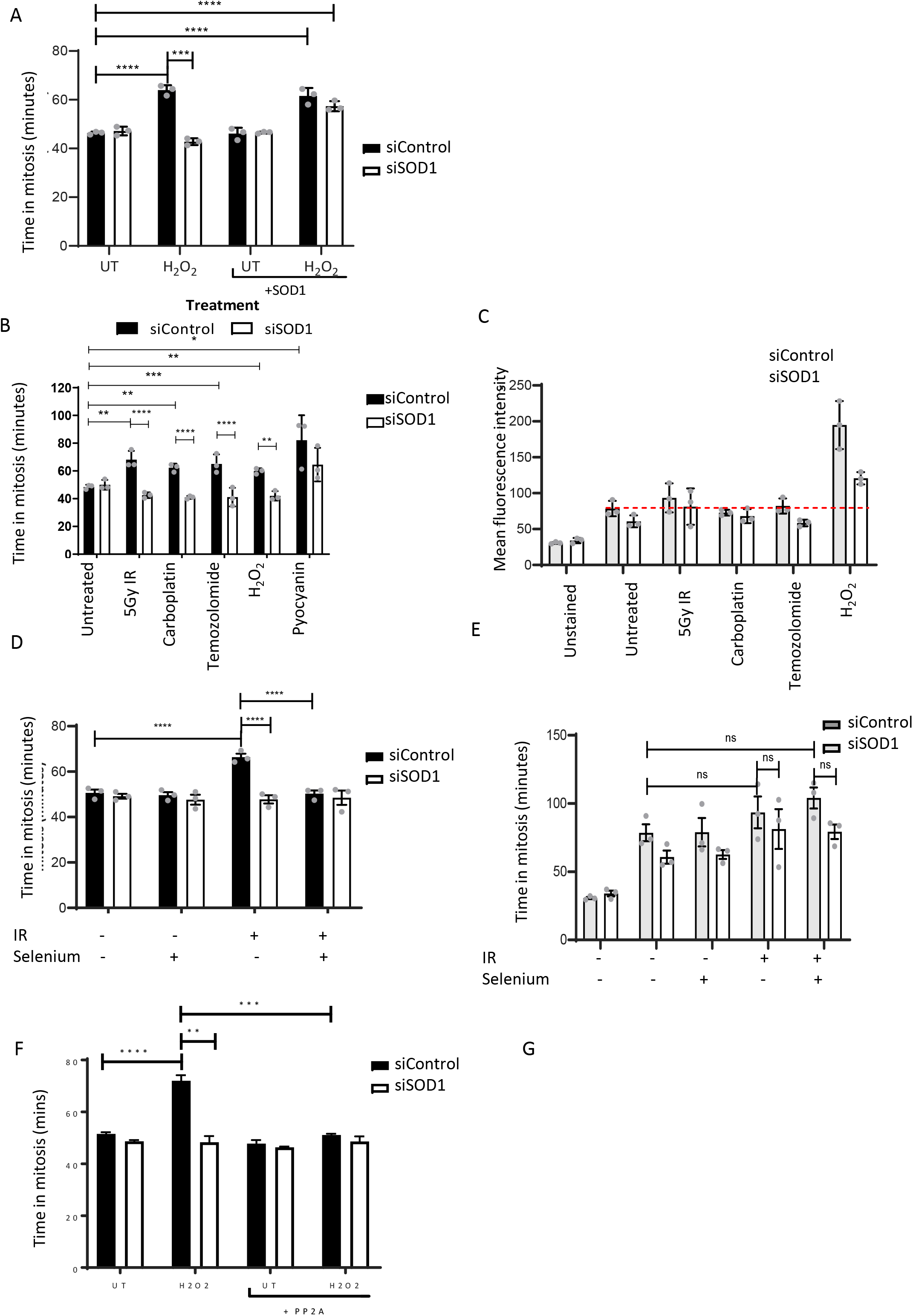
ROS inducing agents also induce SOD1 dependent mitotic arrest. **A, B and D** HeLa cells incubated with the indicated siRNAs were treated as indicated and viewed by time lapse microscopy. Error bars represent mean +/-SEM of 3 independent experiments (n>150). Results were analysed using a oneway ANOVA with Dunnett’s multiple comparison test. *p=0.05, **p=0.01, ***p=0.001, ****p=0.0001 **C and E**. Fluorescent ROS intensity of HeLa cells incubated with the indicated siRNAs and treated as indicated was analysed. Each condition was measured in triplicate, averaged and normalised against the unstained control for each independent repeat +/-SEM (N= 3). Results were analysed using a one-way ANOVA with Dunnett’s multiple comparison test. *p=0.05, **p=0.01, ***p=0.001, ****p=0.0001 **F.** HeLa cells incubated with the indicated siRNAs and expression vectors were treated as indicated and viewed by time lapse microscopy. Error bars represent mean +/-SEM of 3 independent experiments (n>150). Results were analysed using a one-way ANOVA with Dunnett’s multiple comparison test. *p=0.05, **p=0.01, ***p=0.001, ****p=0.0001

Glutathione peroxidase (GPx1) is a ROS-reducing protein which catalyses the reduction of H_2_O_2_ to water and oxygen. Selenium supplementation has been shown to upregulate GPx1. We found that supplementing cells with 50nM selenium led to a detectable increase in GPx1 expression (**Figure S2B and C**) and prevented SOD-dependent, IR-induced mitotic arrest (**Figure 2D**). Surprisingly, selenium supplementation did not reduce cellular ROS (**Figure 2E**) further supporting the hypothesis that H_2_O_2_ is not the cause of the arrest. Furthermore, selenium supplementation also protected against SOD1-dependent mitotic arrest induced by non-ROS-inducing DNA damaging agents (**Figure S2D and 2E**), indicating that selenium prevents mitotic arrest via another pathway. Glutathione metabolism has been implicated in regulation of Protein phosphatase 2A (PP2A), which is required for mitotic exit [27]. Glutathione reduces oxidised PP2A, recovering phosphatase activity [28] and selenium has also been shown to potently stimulate PP2A activity directly [29]. We found that overexpression of PP2A was also able to override DNA damage induced arrest (**Figure 2F**), suggesting one possible mechanism via which selenium overrides the arrest. Taken together, our data suggest that changes in intracellular ROS in the absence of SOD1 do not account for the mitotic delay observed following treatment with DNA damaging agents.

### SOD1 depletion leads to elevated mitotic defects and increased DNA damage but does not impact the SAC

DNA damage induces many aberrant mitotic phenotypes such as micronuclei, DNA bridges, lagging chromosomes and failed cytokinesis (**Figure S3A**), all of which may influence mitotic transit time. We observed all these phenotypes following treatment with IR, however SOD1 depletion induced an additional increase in aberrant mitotic phenotypes (**Figure 3A and S3B**) indicating that SOD1-induced mitotic delay is not a consequence of these phenotypes. Mikhailov *et al* suggest that DNA damage induced mitotic arrest is due to damaged centromeres preventing the SAC from being satisfied [3]. However, whilst we observe damaged centromeres in mitotic cells (as represented by γH2AX) 4Dfollowing treatment with IR, these are not only present but are increased in the absence of SOD1 (**Figure 3B**), indicating that they are not directly responsible for the observed arrest.

**Figure 3.**
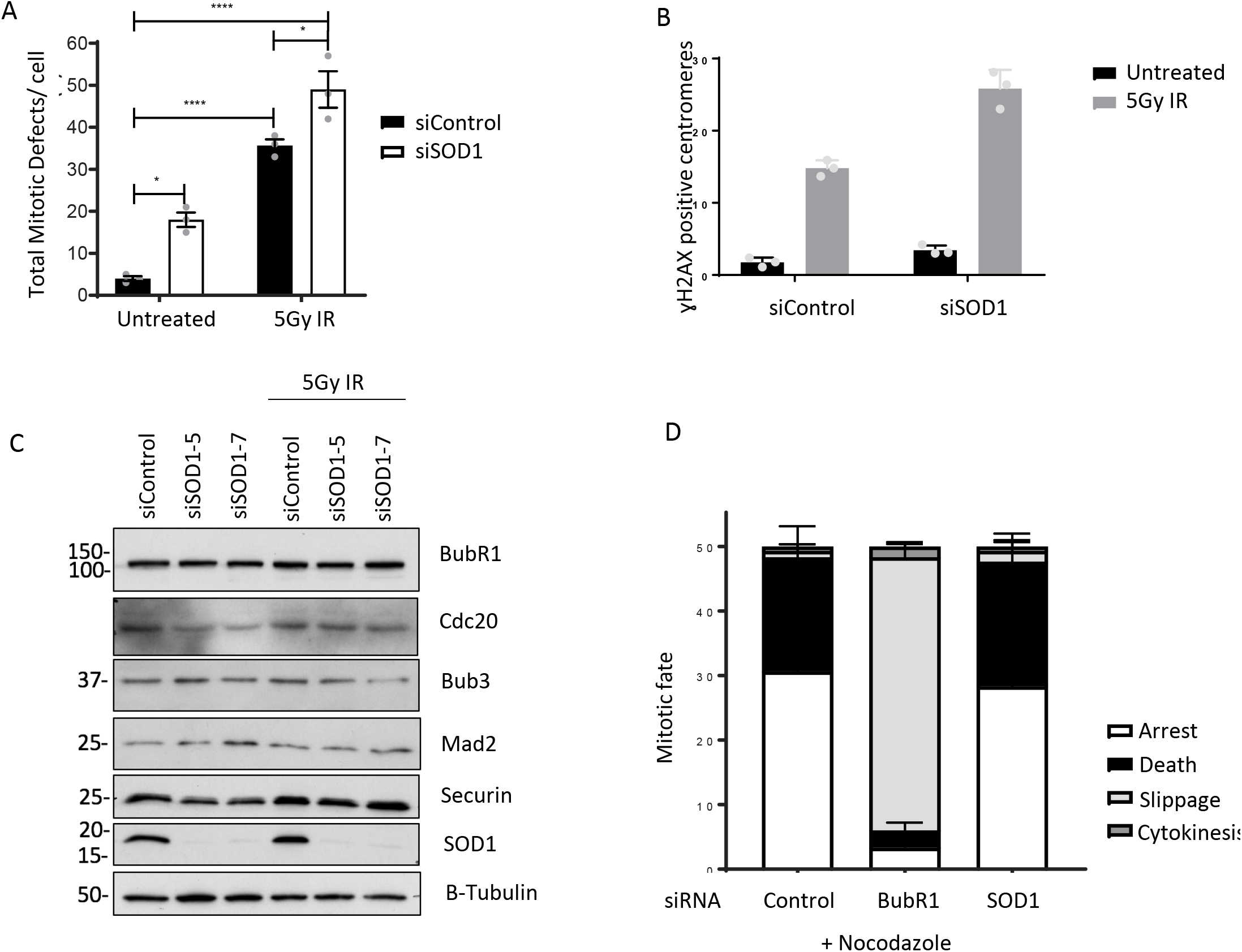
SOD1 depletion leads to increased mitotic defects but leaves the SAC intact. **A.** HeLa cells incubated with the indicated siRNA and treated as indicated were stained with DAPI and visualised by fluorescence microscopy. Mean +/-SEM of 3 independent experiments (n>150). Results were analysed using a one-way ANOVA with Dunnett’s multiple comparison test. *p=0.05, **p=0.01, ***p=0.001, ****p=0.0001 **B.**HeLa cells incubated with the indicated siRNAs and treated as indicated were stained for CenpB and γH2AX. Mean +/-SEM of 3 independent experiments (n=150). Results were analysed using a one-way ANOVA with Dunnett’s multiple comparison test. *p=0.05, **p=0.01, ***p=0.001, ****p=0.0001 **C**. HeLa cells incubated with the indicated siRNAs and treated as indicated were lysed and protein extracts analysed by western blot. **D.**Cells were transfected with indicated siRNAs for 72 hours and treated with nocodazole for the remaining 16 hours. Cells were then visualised by live cell imaging and cell fate was scored.

In lower eukaryotes, DNA damage induces mitotic arrest dependent on the spindle assembly checkpoint (SAC) [30]. To determine whether SOD1 as both a catalytic enzyme and a transcription factor to control the SAC, we first studied the protein and mRNA levels of the mitotic checkpoint complex (MCC) which are required for SAC arrest. qPCR revealed no significant transcriptional changes of the SAC proteins following SOD1 siRNA depletion (**Figure S4A**) and this also translated to the protein level (**Figure 3C**). To confirm that SOD1 does not affect the SAC, we treated cells with the mitotic inhibitor nocodazole, which leads to microtubule destabilisation and permanent SAC activation. When treated with nocodazole, cells arrest in mitosis at the SAC and do not undergo cytokinesis. BUBR1 (an essential protein in the SAC) depletion abrogates nocodazole-induced arrest and the majority of cells escape mitosis without dividing (a process known as “mitotic slippage”). However, following SOD1 depletion, we observed no significant change in the proportion of arrest, death or slippage when compared with the control sample, indicating that SOD1 is not required for initiation or maintenance of the SAC (**Figure 3D**).

### SOD1 is involved in the DDR

SOD1 mutations lead to the accumulation of DNA damage [20] indicating a role for SOD1 in the DDR. SOD1 accumulates in the nucleus in response to H_2_O_2_ [23,25]. We show that this is also the case in response to IR (**Figure 4A,B**) We also observed higher levels of γH2AX foci, a DNA damage marker, in SOD1-depleted cells, than in cell transfected with a non-targeting control siRNA cells following treatment with H_2_O_2_ or exposure to IR **Figure 4C**). Increased DNA damage in SOD1-depleted cells was further confirmed via COMET assay (**Figure 4D and E**). However, despite higher or equal levels of damage (**Figure 4F)**, SOD1 depletion resulted in a reduction in observable RAD51 foci 4 hrs post-IR, indicating reduced DNA repair capacity (**Figure 4G**).

**Figure 4.**
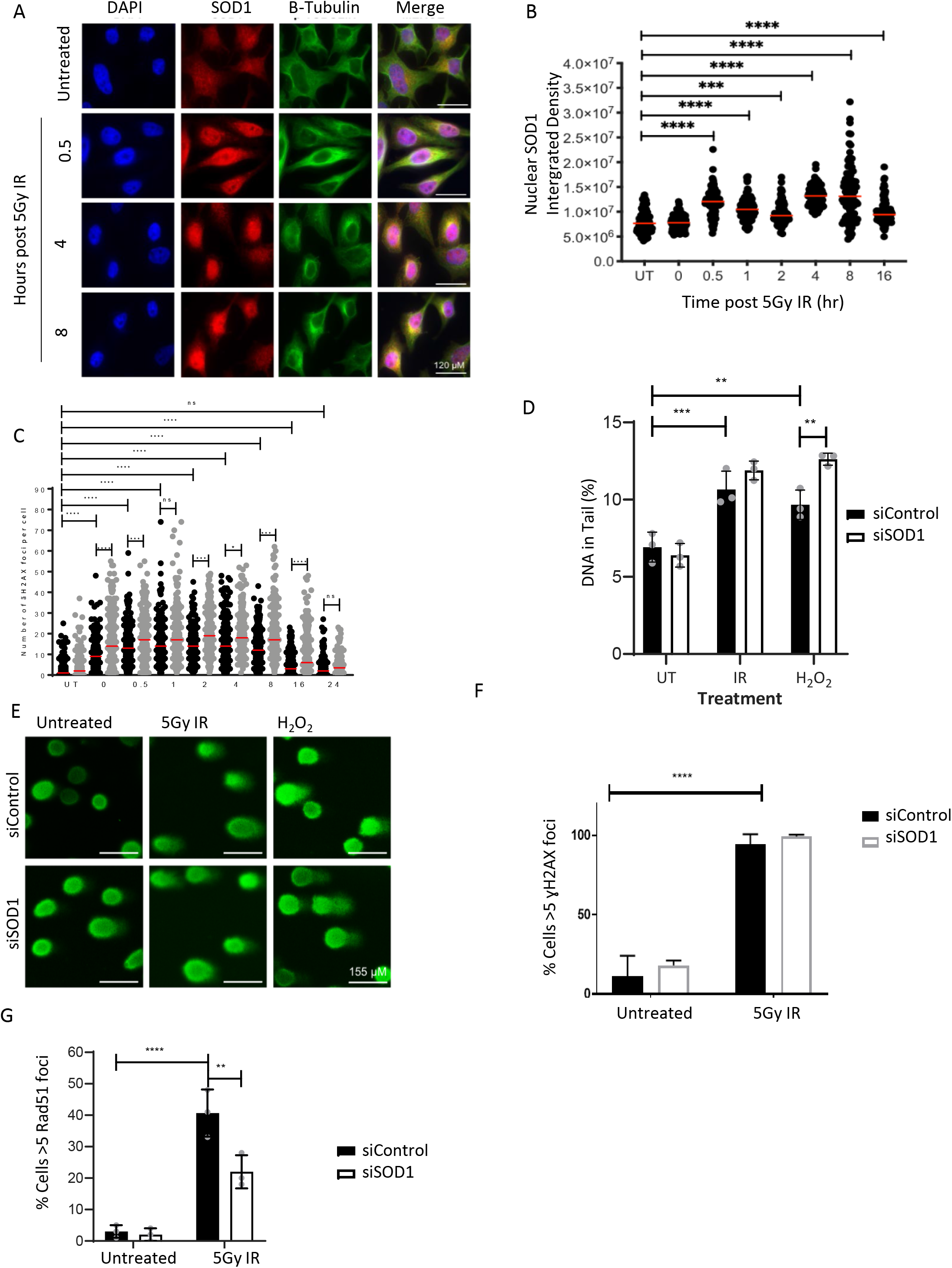
SOD1 is involved in the DDR. **A.** HeLa cells incubated with the indicated siRNA and treated as indicated were stained for SOD1 and viewed by fluorescence microscopy**. B.** Analysis of A using image J to calculate relative fluorescent intensity of SOD1 in DAPI positive areas. **C.** HeLa cells incubated with the siControl (Black) and siSOD1 (Grey) were treated with 5Gy IR and incubated as indicated and stained for γH2AX were viewed by fluorescence microscopy and number of γH2AX foci was scored. Collective data from 3 individual experiments (n=150) Results were analysed using a one-way ANOVA with Dunnett’s multiple comparison test. *p=0.05, **p=0.01, ***p=0.001, ****p=0.0001 **D.** Comet assay was performed on HeLa cells incubated with the indicated siRNA and treated as indicated. % DNA in tail was calculated using cometscore software. Mean +/-SEM of 3 independent experiments (n>15O). Results were analysed using a one-way ANOVA with Dunnett’s multiple comparison test. *p=0.05, **p=0.01, ***p=0.001, ****p=0.0001. **E**. Representative images from **D**. **F.**HeLa cells incubated with the indicated siRNA and treated as indicated were stained for γH2AX were viewed by fluorescence microscopy. Mean =/-SEM of 3 independent experiments (n>150). **G**. HeLa cells incubated with the indicated siRNA and treated as indicated were stained for Rad51 were viewed by fluorescence microscopy. Mean =/-SEM of 3 independent experiments (n>150). Results were analysed using a one-way ANOVA with Dunnett’s multiple comparison test. *p=0.05, **p=0.01, ***p=0.001, ****p=0.0001

### SOD1 depletion leads to reduced Mitotic EdU incorporation

Limited DNA replication and repair have been observed in mitosis via incorporation of EdU in response to replication stress [15,31,32] and more recently, DNA damage [17]. We observed that SOD1 promotes efficient mitotic EdU incorporation in response to both replication stress (**Figure 5A,B**) and DNA damage (**Figure 5C**). Rad51 [16] and Rad52 [31] have both been demonstrated to play a role in mitotic EdU incorporation. Our data demonstrates that SOD1 depletion reduces mitotic EdU incorporation to an extent similar to the RAD51 inhibitor (BO2) and this reduction was not synergistic in combination with BO2 treatment (**Figure 5d).**RAD52 inhibition did not reduce the levels of mitotic EdU incorporation to the same extent as SOD1 depletion or RAD51 inhibition but displayed an additive decrease in combination with SOD1 inhibition (**Figure 5d**). Whilst the RAD51 inhibitor led to complete mitotic arrest and cell death independently of DNA damage meaning data collection was not possible, the RAD52 inhibitor restored mitotic arrest in SOD1-depleted cells (**Figure 5e**) in response to DNA damage, implying that the defects in mitotic delay induced by SOD1 loss are driven by RAD52-dependent repair pathways.

**Figure 5.**
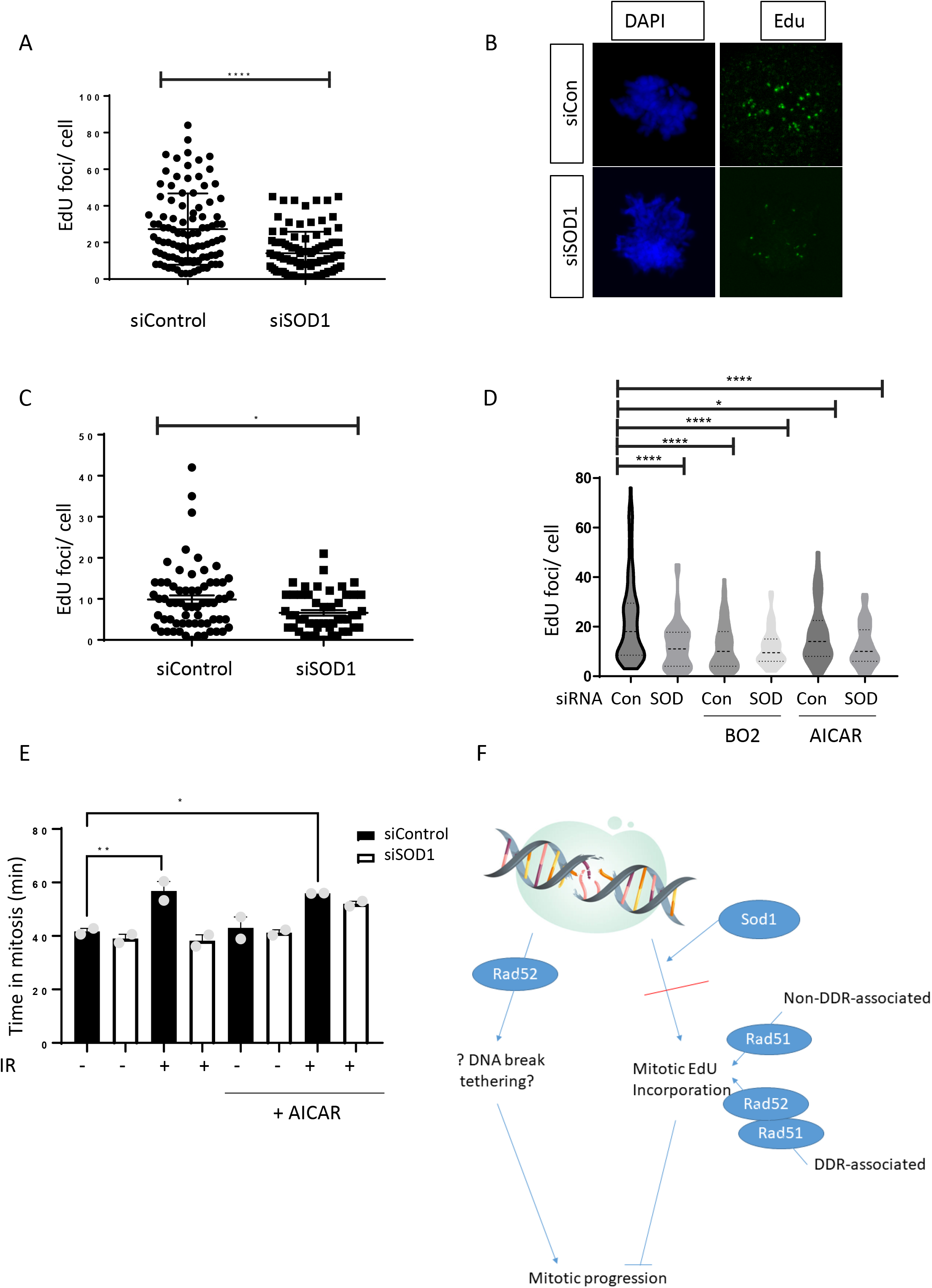
SOD1 promotes effective mitotic EdU incorporation in a Rad51 and Rad52 independent manner. **A.** HeLa cells were incubated with the indicated siRNAs 48 hours prior to treatment with 0.4μM aphidicolin. Following aphidicolin, cells were release into 9μM RO3306 for 5 hours to arrest in G2. Cells were released into mitosis and pulsed for 30 minutes with EdU. Number of EdU foci per mitotic cell was scored. Error bars represent SEM of 3 individual repeats. N=150. Results were analysed using a one-way ANOVA with Dunnett’s multiple comparison test. *p=0.05, **p=0.01, ***p=0.001, ****p=0.0001 **B.** Representative images of A. **C.**HeLa cells were incubated with the indicated siRNAs 48 hours prior to treatment with 5Gy IR, followed by G2 arrest with 9μM RO3306 1 hour later for 5 hours. Cells were released into mitosis and then pulsed for 30 minutes with EdU. Number of EdU foci per mitotic cell was scored. Error bars represent SEM of 3 individual repeats. N=150. Results were analysed using a one-way ANOVA with Dunnett’s multiple comparison test. *p=0.05, **p=0.01, ***p=0.001, ****p=0.0001 **D.**As in A. In the case of chemical inhibition, cells were pulsed for 30 minutes with EdU in the presence of either 20μM B02 or 20μM AICAR. Results were analysed using a one-way ANOVA with Dunnett’s multiple comparison test. *p=0.05, **p=0.01, ***p=0.001, ****p=0.0001 **E.** HeLa cells were incubated with the indicated siRNA for 48 hours prior to treatment with 5Gy IR and Rad52 inhibitor. Cells were anlysed by time lapse microscopy for 20 hours. Error bars represent mean and SEM of 3 independent biological replicates. N=150. Results were analysed using a one-way ANOVA with Dunnett’s multiple comparison test. *p=0.05, **p=0.01, ***p=0.001, ****p=0.0001 **F.**Proposed model.

## Discussion

Here we demonstrate a novel role for Superoxide Dismutase 1 in control of the cell cycle and DNA repair following induction of DNA damage. We have uncovered a SOD1-dependent mechanism that drives mitotic slowing in the presence of DNA damage. In the absence of this delay, we observe elevated levels of DNA damage, increased mitotic defects and reduced mitotic DNA synthesis.

Previous reports have proposed that damage to kinetochores causes delayed mitotic progression though persistent SAC activation [3,16]. However, we found that whilst cells treated with IR exhibit high levels of damaged centromeres, this was elevated further in cells depleted of SOD1. Since SOD1-depleted cells do not exhibit DNA damage-induced mitotic arrest but are capable of nocodazole-induced mitotic arrest we propose that DNA damage leads to prolonged mitotic progression through a pathway distinct from the canonical SAC.

Previously studies suggest that RAD51 inhibition leads to reduced mitotic EdU incorporation and prolonged mitosis and it was hypothesized that mitotic exit cannot occur until mitotic DNA synthesis is complete [16]. Surprisingly, we observed that whilst SOD1 depletion has the opposite effect to RAD51 inhibition in that it promotes mitotic exit, SOD1 depletion phenocopied Rad51 inhibition and reduced mitotic EdU incorporation.RAD52 has been shown to be important for mitotic EdU incorporation [31] though to a lesser extent than RAD51 [16] and our data reflect this.However, whilst SOD1 depletion had no effect on mitotic EdU incorporation in the absence of RAD51, SOD1 depletion led to a further reduction in mitotic EdU incorporation in the absence of RAD52. This can potentially be explained by the two types of mitotic EdU incorporation described by Wassing et al [16]; RAD51-dependent non-DDR-associated MiDAS which occurs in the absence of breaks and RAD52-dependent DDR-associated MiDAS which occurs in the presence of breaks. We propose that SOD1 and Rad51 are required for both pathways, but Rad52 is required only for DDR-associated MiDAS.

Interestingly, we found that mitotic progression in the absence of SOD1 is dependent on RAD52. We propose that there are two potential fates for mitotic DNA breaks; mitotic delay and repair through MiDAS or mitotic progression, resulting in mitotic defects, followed by repair in G1. Our model (**Figure 5F)**suggests the existence of a fine balance between these two fates, controlled in part by SOD1 and RAD52. We propose that in response to DNA breaks, SOD1 guides cells down a pathway whereby mitosis is slowed and mitotic DNA synthesis and repair is instigated, however in the absence of SOD1, this pathway can no longer function, meaning cells prepare for repair in the subsequent G1. We propose a “point of no return” in this pathway, after SOD1 but before RAD51. If cells are guided down the repair in mitosis pathway by a functional SOD1 but RAD51 is not present, a prolonged mitosis results but repair cannot be completed resulting in mitotic cell death.

In this study, we demonstrate the existence of a signaling cascade leading to prolonged mitosis and mitotic DNA synthesis following genotoxic stress induced by DNA damage or replication defects.

Further work is required to elucidate the mechanism controlling mitotic progression in the presence of DNA damage and whether this involves SAC components-however, our data indicate that the canonical SAC is active in the absence of SOD1. These data suggest the existence of a previously unknown DNA damage checkpoint in mitosis which detects DNA damage and prolongs mitosis to allow for mitotic DNA repair. This checkpoint appears to be independent of the interphase DNA damage checkpoints and could be a potential druggable target for future therapeutics design.

## Supplemental Figures

**Figure S1A.**
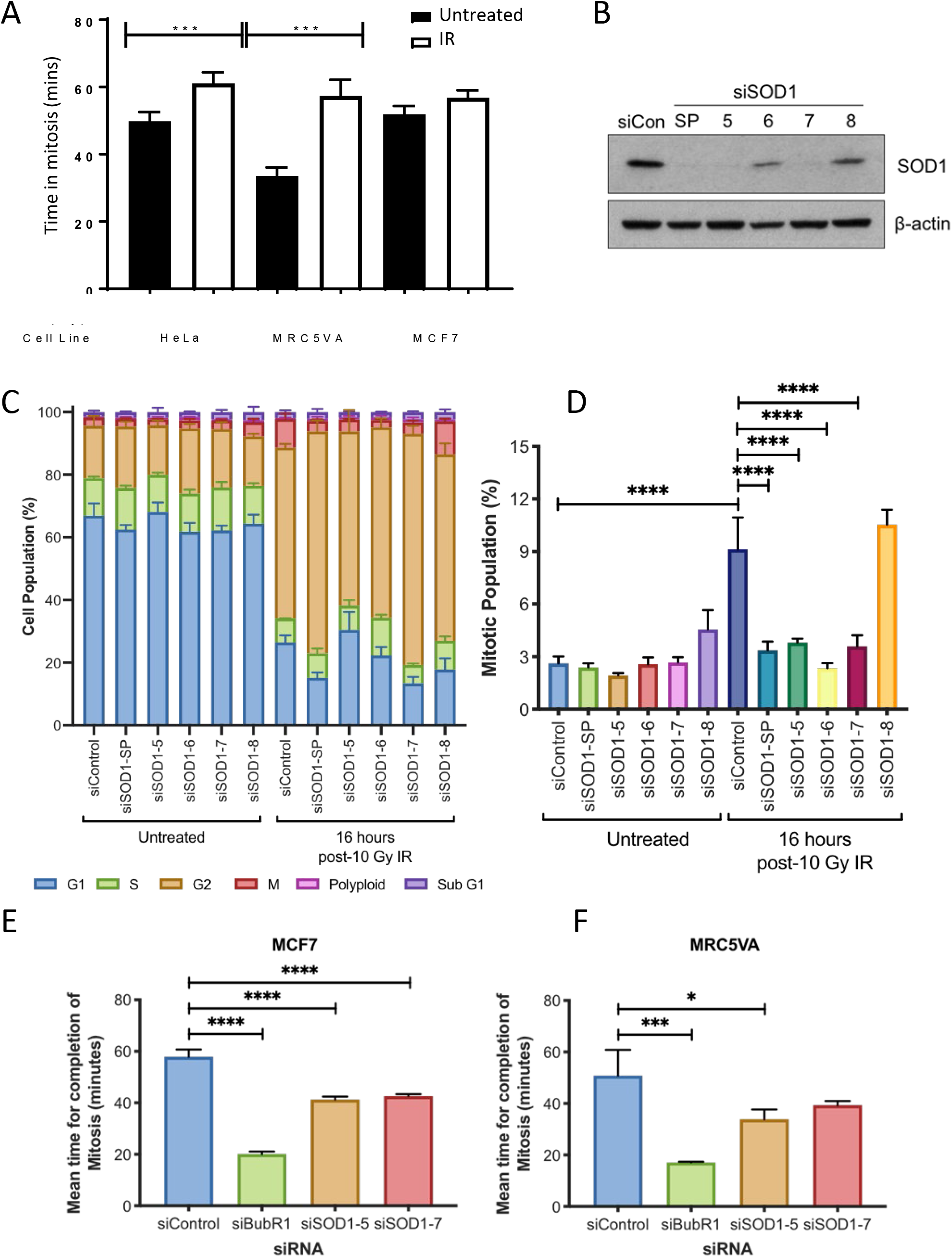
HeLa, MRC5VA and MCF7 cells were treated as indicated and analysed by time lapse microscopy for 16 hours. B. HeLa cells were incubated with the indicated siRNAs for 48 hours and analysed by western blotting. C,D. HeLa cells were incubated for 48 hours with the indicated siRNAs, stained with propidium iodide and pH3-FITC and analysed by flow cytometry. E. MCF7 and F. MRC5VA cells were treated as indicated and analysed by time;apse microscopy.

**Figure S2A.**
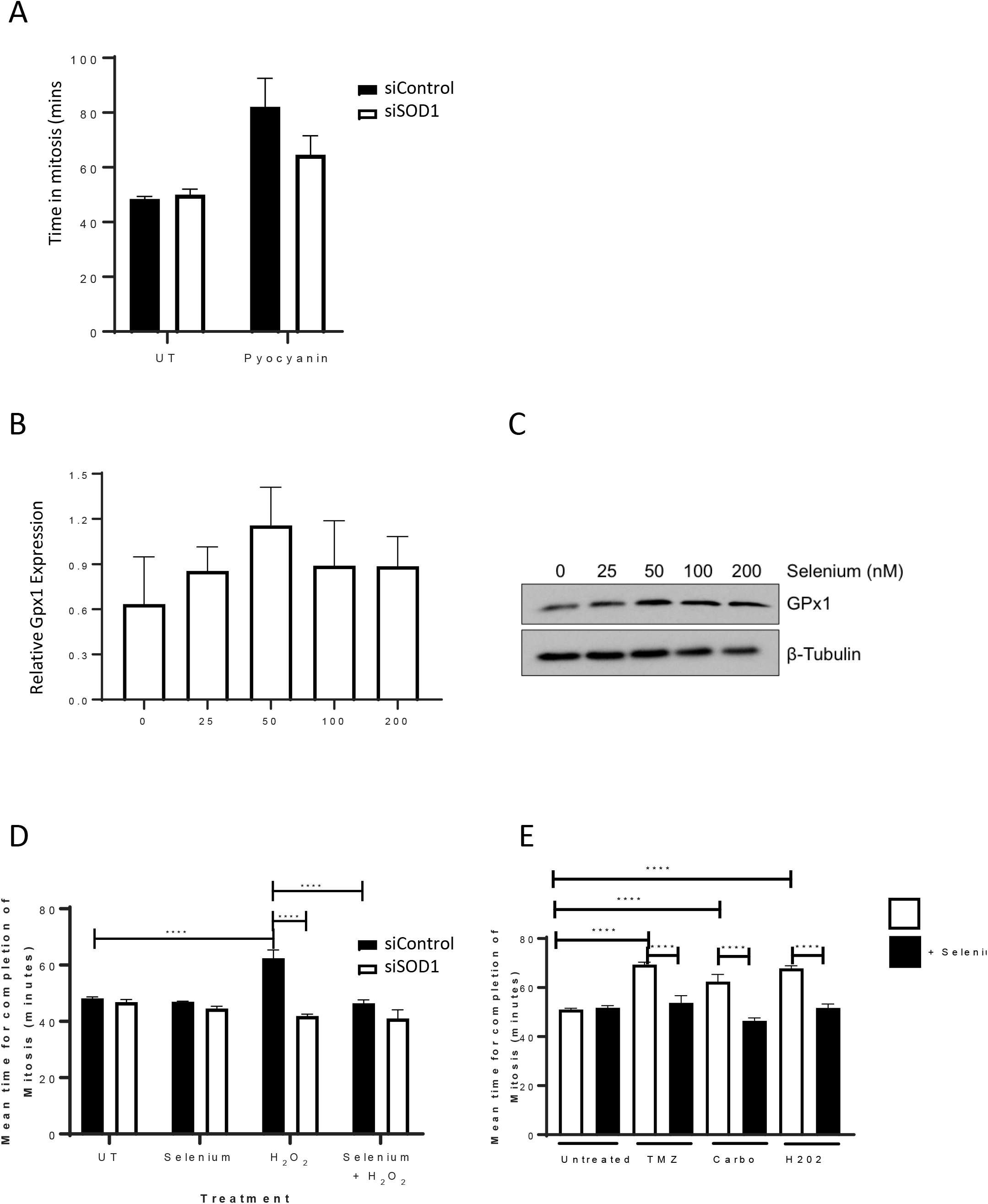
HeLa cells were incubated with the indicated siRNAs for 48 hours and treated as indicated prior to analysis by time lapse microscopy for 16 hours post treatment. B. HeLa cells were incubated with sodium selenite (nM) for 48 hours and analysed by western blotting. Densitometry performed on 3 individual repeats. C. Representative blot from B. D,E HeLa cells were treated as indicated and analysed by time lapse microscopy for 16 hours post treatment.

**Figure S3A.**
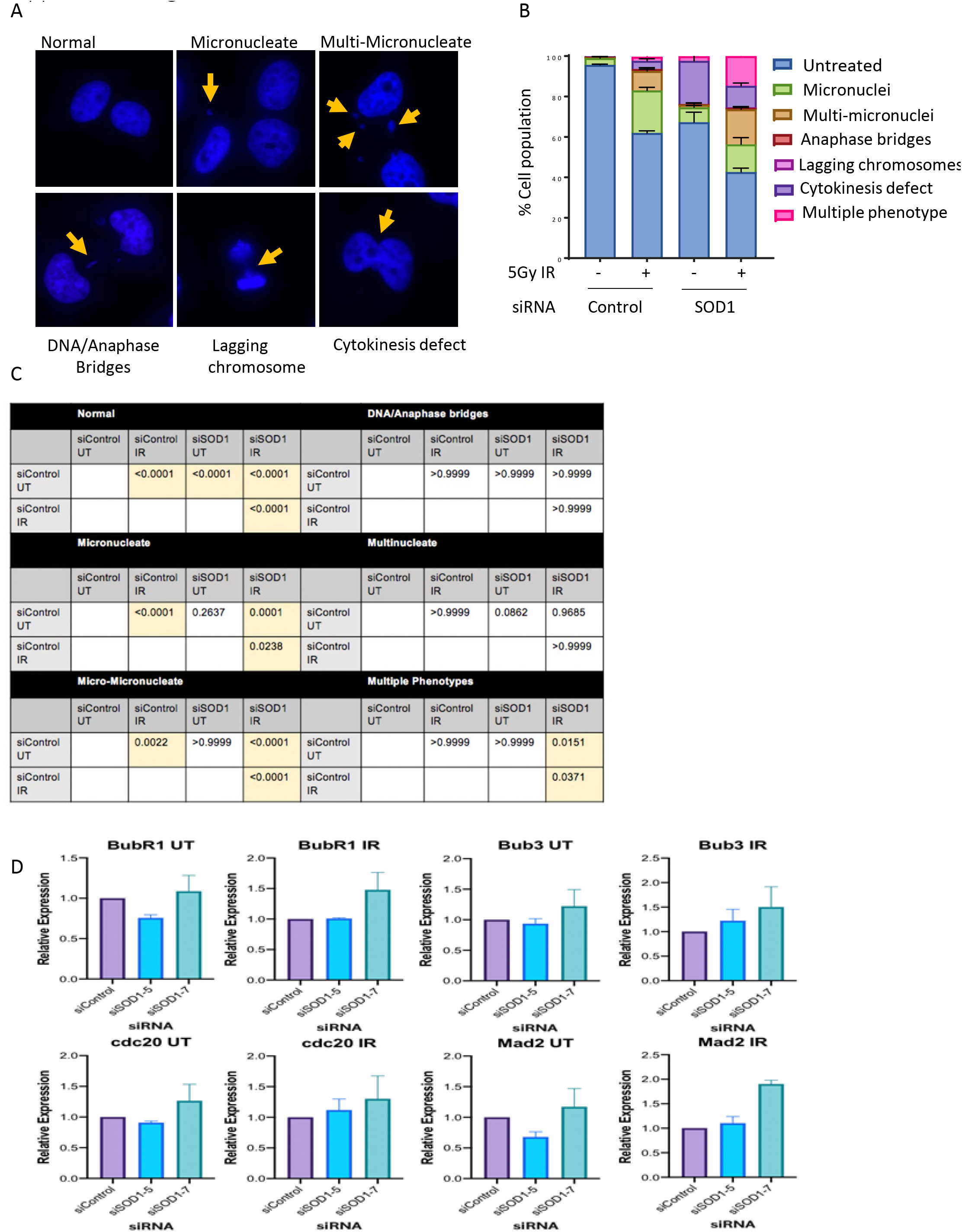
Representative images of B. HeLa cells were treated as indicated, stained with DAPI and analysed by fluorescent microscopy. C Statistical analysis of B. D. HeLa cells were treated as indicated and RNA was extracted and analysed by qRT-PCR for MCC component expression.

## References

1. Bartek, J.; Lukas, C.; Lukas, J. Checking on DNA damage in S phase. Nat Rev Mol Cell Biol 2004, 5, 792–804, doi:10.1038/nrm1493.

2. Smits, V.A.; Klompmaker, R.; Arnaud, L.; Rijksen, G.; Nigg, E.A.; Medema, R.H. Polo-like kinase-1 is a target of the DNA damage checkpoint. Nat Cell Biol 2000, 2, 672–676, doi:10.1038/35023629.

3. Mikhailov, A.; Cole, R.W.; Rieder, C.L. DNA damage during mitosis in human cells delays the metaphase/anaphase transition via the spindle-assembly checkpoint. Curr Biol 2002, 12, 1797–1806.

4. Rello-Varona, S.; Gamez, A.; Moreno, V.; Stockert, J.C.; Cristobal, J.; Pacheco, M.; Canete, M.; Juarranz, A.; Villanueva, A. Metaphase arrest and cell death induced by etoposide on HeLa cells. Int J Biochem Cell Biol 2006, 38, 2183–2195, doi:10.1016/j.biocel.2006.06.013.

5. Thompson, R.; Shah, R.B.; Liu, P.H.; Gupta, Y.K.; Ando, K.; Aggarwal, A.K.; Sidi, S. An Inhibitor of PIDDosome Formation. Mol Cell 2015, 58, 767–779, doi:10.1016/j.molcel.2015.03.034.

6. Blackford, A.N.; Stucki, M. How Cells Respond to DNA Breaks in Mitosis. Trends Biochem Sci 2020, 45, 321–331, doi:10.1016/j.tibs.2019.12.010.

7. Kastan, M.B.; Bartek, J. Cell-cycle checkpoints and cancer. Nature 2004, 432, 316–323, doi:10.1038/nature03097.

8. Orthwein, A.; Fradet-Turcotte, A.; Noordermeer, S.M.; Canny, M.D.; Brun, C.M.; Strecker, J.; Escribano-Diaz, C.; Durocher, D. Mitosis inhibits DNA double-strand break repair to guard against telomere fusions. Science 2014, 344, 189–193, doi:10.1126/science.1248024.

9. Giunta, S.; Belotserkovskaya, R.; Jackson, S.P. DNA damage signaling in response to doublestrand breaks during mitosis. J Cell Biol 2010, 190, 197–207, doi:10.1083/jcb.200911156.

10. Royou, A.; Gagou, M.E.; Karess, R.; Sullivan, W. BubR1- and Polo-coated DNA tethers facilitate poleward segregation of acentric chromatids. Cell 2010, 140, 235–245, doi:10.1016/j.cell.2009.12.043.

11. Leimbacher, P.A.; Jones, S.E.; Shorrocks, A.K.; de Marco Zompit, M.; Day, M.; Blaauwendraad, J.; Bundschuh, D.; Bonham, S.; Fischer, R.; Fink, D., et al. MDC1 Interacts with TOPBP1 to Maintain Chromosomal Stability during Mitosis. Mol Cell 2019, 10.1016/j.molcel.2019.02.014, doi:10.1016/j.molcel.2019.02.014.

12. Burrow, A.A.; Williams, L.E.; Pierce, L.C.; Wang, Y.H. Over half of breakpoints in gene pairs involved in cancer-specific recurrent translocations are mapped to human chromosomal fragile sites. BMC Genomics 2009, 10, 59, doi:10.1186/1471-2164-10-59.

13. Burrell, R.A.; McClelland, S.E.; Endesfelder, D.; Groth, P.; Weller, M.C.; Shaikh, N.; Domingo, E.; Kanu, N.; Dewhurst, S.M.; Gronroos, E., et al. Replication stress links structural and numerical cancer chromosomal instability. Nature 2013, 494, 492–496, doi:10.1038/nature11935.

14. Le Beau, M.M.; Rassool, F.V.; Neilly, M.E.; Espinosa, R., 3rd; Glover, T.W.; Smith, D.I.; McKeithan, T.W. Replication of a common fragile site, FRA3B, occurs late in S phase and is delayed further upon induction: implications for the mechanism of fragile site induction. Hum Mol Genet 1998, 7, 755–761, doi:10.1093/hmg/7.4.755.

15. Minocherhomji, S.; Ying, S.; Bjerregaard, V.A.; Bursomanno, S.; Aleliunaite, A.; Wu, W.; Mankouri, H.W.; Shen, H.; Liu, Y.; Hickson, I.D. Replication stress activates DNA repair synthesis in mitosis. Nature 2015, 528, 286–290, doi:10.1038/nature16139.

16. Wassing, I.E.; Graham, E.; Saayman, X.; Rampazzo, L.; Ralf, C.; Bassett, A.; Esashi, F. The RAD51 recombinase protects mitotic chromatin in human cells. Nat Commun 2021, 12, 5380, doi:10.1038/s41467-021-25643-y.

17. Gomez Godinez, V.; Kabbara, S.; Sherman, A.; Wu, T.; Cohen, S.; Kong, X.; Maravillas-Montero, J.L.; Shi, Z.; Preece, D.; Yokomori, K., et al. DNA damage induced during mitosis undergoes DNA repair synthesis. PLoS One 2020, 15, e0227849, doi:10.1371/journal.pone.0227849.

18. McCord, J.M.; Fridovich, I. Superoxide dismutase. An enzymic function for erythrocuprein (hemocuprein). J Biol Chem 1969, 244, 6049–6055.

19. Barbosa, L.F.; Cerqueira, F.M.; Macedo, A.F.; Garcia, C.C.; Angeli, J.P.; Schumacher, R.I.; Sogayar, M.C.; Augusto, O.; Carri, M.T.; Di Mascio, P., et al. Increased SOD1 association with chromatin, DNA damage, p53 activation, and apoptosis in a cellular model of SOD1-linked ALS. Biochim Biophys Acta 2010, 1802, 462–471, doi:10.1016/j.bbadis.2010.01.011.

20. Sau, D.; De Biasi, S.; Vitellaro-Zuccarello, L.; Riso, P.; Guarnieri, S.; Porrini, M.; Simeoni, S.; Crippa, V.; Onesto, E.; Palazzolo, I., et al. Mutation of SOD1 in ALS: a gain of a loss of function. Hum Mol Genet 2007, 16, 1604–1618, doi:10.1093/hmg/ddm110.

21. Wang, X.D.; Zhu, M.W.; Shan, D.; Wang, S.Y.; Yin, X.; Yang, Y.Q.; Wang, T.H.; Zhang, C.T.; Wang, Y.; Liang, W.W., et al. Spy1, a unique cell cycle regulator, alters viability in ALS motor neurons and cell lines in response to mutant SOD1-induced DNA damage. DNA Repair (Amst) 2019, 74, 51–62, doi:10.1016/j.dnarep.2018.12.005.

22. Carter, C.D.; Kitchen, L.E.; Au, W.C.; Babic, C.M.; Basrai, M.A. Loss of SOD1 and LYS7 sensitizes Saccharomyces cerevisiae to hydroxyurea and DNA damage agents and downregulates MEC1 pathway effectors. Mol Cell Biol 2005, 25, 10273–10285, doi:10.1128/MCB.25.23.10273-10285.2005.

23. Tsang, C.K.; Liu, Y.; Thomas, J.; Zhang, Y.J.; Zheng, X.F.S. Superoxide dismutase 1 acts as a nuclear transcription factor to regulate oxidative stress resistance. Nat Commun 2014, 5, doi:10.1038/ncomms4446.

24. Bordoni, M.; Pansarasa, O.; Dell’Orco, M.; Crippa, V.; Gagliardi, S.; Sproviero, D.; Bernuzzi, S.; Diamanti, L.; Ceroni, M.; Tedeschi, G., et al. Nuclear Phospho-SOD1 Protects DNA from Oxidative Stress Damage in Amyotrophic Lateral Sclerosis. J Clin Med 2019, 8, doi:10.3390/jcm8050729.

25. Li, X.; Qiu, S.; Shi, J.; Wang, S.; Wang, M.; Xu, Y.; Nie, Z.; Liu, C.; Liu, C. A new function of copper zinc superoxide dismutase: as a regulatory DNA-binding protein in gene expression in response to intracellular hydrogen peroxide. Nucleic Acids Res 2019, 47, 5074–5085, doi:10.1093/nar/gkz256.

26. Garribba, L., Wu, L., Ozer, O., Bhowmick, R., Hickson, ID. Inducing and Detecting Mitotic DNA Synthesis at Difficult-to-Replicate Loci. In Methods in Enzymology, Elsevier: 2018; Vol. 601.

27. Forester, C.M.; Maddox, J.; Louis, J.V.; Goris, J.; Virshup, D.M. Control of mitotic exit by PP2A regulation of Cdc25C and Cdk1. Proc Natl Acad Sci U S A 2007, 104, 19867–19872, doi:10.1073/pnas.0709879104.

28. Rao, R.K.; Clayton, L.W. Regulation of protein phosphatase 2A by hydrogen peroxide and glutathionylation. Biochem Biophys Res Commun 2002, 293, 610–616, doi:10.1016/S0006-291X(02)00268-1.

29. Corcoran, N.M.; Martin, D.; Hutter-Paier, B.; Windisch, M.; Nguyen, T.; Nheu, L.; Sundstrom, L.E.; Costello, A.J.; Hovens, C.M. Sodium selenate specifically activates PP2A phosphatase, dephosphorylates tau and reverses memory deficits in an Alzheimer’s disease model. J Clin Neurosci 2010, 17, 1025–1033, doi:10.1016/j.jocn.2010.04.020.

30. Kim, E.M.; Burke, D.J. DNA damage activates the SAC in an ATM/ATR-dependent manner, independently of the kinetochore. PLoS Genet 2008, 4, e1000015, doi:10.1371/journal.pgen.1000015.

31. Bhowmick, R.; Minocherhomji, S.; Hickson, I.D. RAD52 Facilitates Mitotic DNA Synthesis Following Replication Stress. Mol Cell 2016, 64, 1117–1126, doi:10.1016/j.molcel.2016.10.037.

32. Mankouri, H.W.; Huttner, D.; Hickson, I.D. How unfinished business from S-phase affects mitosis and beyond. EMBO J 2013, 32, 2661–2671, doi:10.1038/emboj.2013.211.

